# Diversity and evolution of nitric oxide reduction

**DOI:** 10.1101/2021.10.15.464467

**Authors:** Ranjani Murali, Laura A. Pace, Robert A. Sanford, Lewis M. Ward, Mackenzie Lynes, Roland Hatzenpichler, Usha F. Lingappa, Woodward W. Fischer, Robert B. Gennis, James Hemp

## Abstract

Nitrogen is an essential element for life, with the availability of fixed nitrogen limiting productivity in many ecosystems. The return of oxidized nitrogen species to the atmospheric N_2_ pool is predominately catalyzed by microbial denitrification (NO_3_^-^ → NO_2_^-^ → NO → N_2_O → N_2_)^1^. Incomplete denitrification can produce N_2_O as a terminal product, leading to an increase in atmospheric N_2_O, a potent greenhouse and ozone-depleting gas^2^. The production of N_2_O is catalyzed by nitric oxide reductase (NOR) members of the heme-copper oxidoreductase (HCO) superfamily^3^. Here we use phylogenomics to identify a number of previously uncharacterized HCO families and propose that many of them (eNOR, sNOR, gNOR, and nNOR) perform nitric oxide reduction. These families have novel active-site structures and several have conserved proton channels, suggesting that they might be able to couple nitric oxide reduction to energy conservation. We isolated and biochemically characterized a member of the eNOR family from *Rhodothermus marinus*, verifying that it performs nitric oxide reduction both *in vitro* and *in vivo*. These newly identified NORs exhibit broad phylogenetic and environmental distributions, expanding the diversity of microbes that can perform denitrification. Phylogenetic analyses of the HCO superfamily demonstrate that nitric oxide reductases evolved multiple times independently from oxygen reductases, suggesting that complete denitrification evolved after aerobic respiration.

---

The HCO superfamily is extremely diverse, with members playing crucial roles in both aerobic (oxygen reductases) and anaerobic (nitric oxide reductases) respiration. The superfamily currently consists of at least three oxygen reductase families (A, B and C) and three NOR families (cNOR, qNOR, and qCu_A_NOR)^4^. The oxygen reductases catalyze the reduction of O_2_ to water (O_2_ + 4e_out_^-^ + 4H_in_^+^ + nH_in_^+^ → 2H_2_O + nH_out_^+^) and share a conserved reaction mechanism^5,6^, where three of the electrons required to reduce O_2_ are provided by the active-site metals, heme-Fe and Cu_B_, while the fourth electron is derived from a redox-active cross-linked tyrosine cofactor^7^ (**Figure 1**). The free energy of the reaction is converted to a transmembrane proton electrochemical gradient by two different mechanisms, charge separation across the membrane and proton pumping^8–10^. Both the chemical and pumped protons are taken up from the electrochemically negative side of the membrane (bacterial cytoplasm) via conserved proton-conducting channels that are comprised of conserved polar residues and internal water molecules. The different oxygen reductase families exhibit differential proton pumping stochiometries (n=4 for the A-family, and n=2 for the B and C-families)^8–10^, depending on the details of their conserved proton channels. The oxygen reductases also vary in their secondary subunits that function as redox relays from electron donors to the active-site, with the A and B-families utilizing a Cu_A_-containing subunit^11–13^ and the C-family containing one or more cytochrome *c* subunits^14^ (**Figure 1**).

**Figure 1.**
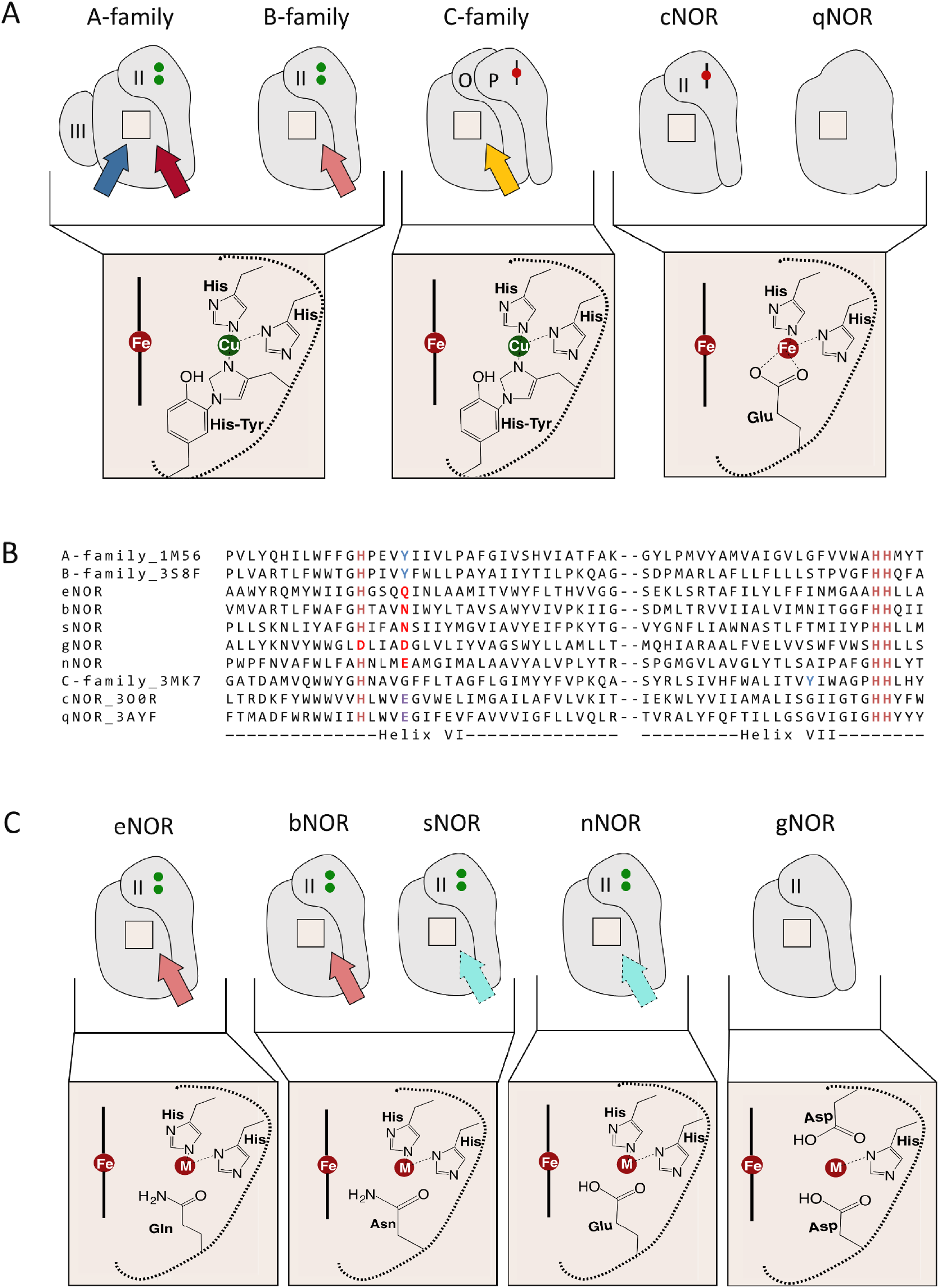
Comparison of HCO active sites. a) Active-site and proton channel properties of the five characterized HCO families (A-family, B-family, C-family, cNOR, and qNOR). The oxygen reductases all have an active-site composed of high-spin heme, a redox-active cross-linked tyrosine cofactor, and a copper (Cu_B_) ligated by three histidines. The A-family has two conserved proton channels, whereas the B and C-families only have one. The active-sites of the nitric oxide reductases are composed of a high-spin heme and an iron (Fe_B_) that is ligated by three histidines and a glutamate. Notably they are missing the tyrosine cofactor. The cNOR and qNOR are also missing conserved proton channels, making them non-electrogenic. b) Sequence alignment of the active-sites of the newly discovered HCO families that are related to the B-family. c) Predicted active-sites and proton channels for the new HCO families. The eNOR, bNOR, sNOR, and nNOR families contain completely conserved proton channels shown here as arrows. The putative proton channel in the bNOR and eNOR families are highly similar to the K-channel from the B-family oxygen reductase and are colored in red. The K-channel in the B-family is also similar to the K-channel in the A-family oxygen reductase which is colored in dark red. The proton channel in the C-family is different from these channels and is marked in yellow. The putative proton channels in sNOR and nNOR are marked in cyan and differentiated from the other channels with a dashed black outline.

Nitric oxide reductases (NORs) catalyze the reduction of nitric oxide to nitrous oxide (2NO + 2H_out_^+^ + 2e_out_^-^ + nH_in_^+^ → N_2_O + H_2_O + nH_in_^+^). Nitric oxide reduction is only a 2-electron reaction, so it does not require the cross-linked tyrosine cofactor for catalysis. There are currently three known HCO NORs, the cNOR, qNOR, and qCu_A_NOR. The cNOR and qNOR families have a four amino acid coordinated Fe_B_ ion in their active-sites, in contrast to the three amino acid coordinated Cu_B_ found in oxygen reductases^15,16^. The cNOR and qNOR families are closely related to the C-family oxygen reductases and likely evolved from this oxygen reductase family. In accordance with this relationship, the cNORs have a secondary cytochrome *c*-containing subunit, while in the qNORs the two subunits corresponding to those in the cNORs have been fused to a single subunit that lacks the heme *c* binding. Although, the qNOR from *N. meningitidis*^16^ is proposed to take up protons from the cytoplasm for NO reduction, neither the cNOR nor qNOR have conserved residues that could form a proton channel from the cytoplasm, therefore it is unlikely that they conserve energy via either charge separation or proton pumping (n=0 for the cNOR and qNOR families). Phylogenomic analysis shows that the qCu_A_NOR from *Bacillus azotoformans*^17,18^ is unrelated to the qNOR family, and has been reclassified here as the bNOR family. The bNOR active-site structure is fundamentally different than those from the cNOR and qNOR families **(Figure 1)**. It also contains a conserved proton channel that is very similar to that found in the B-family oxygen reductases (**Figure 1)**, and has been shown to be electrogenic^18^. This has important consequences for the efficiency of energy conservation associated with denitrification^19^.

## Novel heme-copper oxygen reductase homologs

Phylogenomic analyses of genomic and metagenomic data have identified at least six new families belonging to the HCO superfamily (**Figure 2**). All of these families are missing the active-site tyrosine, suggesting that they do not catalyze O_2_ reduction. Furthermore, their active-sites exhibit structural features never seen before within the superfamily (**Figure 1**). One of these families is closely related to qNOR and has been proposed to be a nitric oxide dismutase contributing to O_2_ production in *‘Candidatus* Methylomirabilis oxyfera’^20^. Another family is closely related to cNOR and might be a sulfide and acetylene insensitive nitrous oxide reductase^21,22^. The remaining four families (eNOR, sNOR, nNOR and gNOR) are closely related to the B-family of oxygen reductases (**Figure 2**) and encode for homologs of the Cu_A-_containing secondary subunits, consistent with this evolutionary relationship (**Figure 1, Table S1**). Based on modeled active-site structures and genomic context we propose that these four families perform nitric oxide reduction (**Figure 1**).

**Figure 2.**
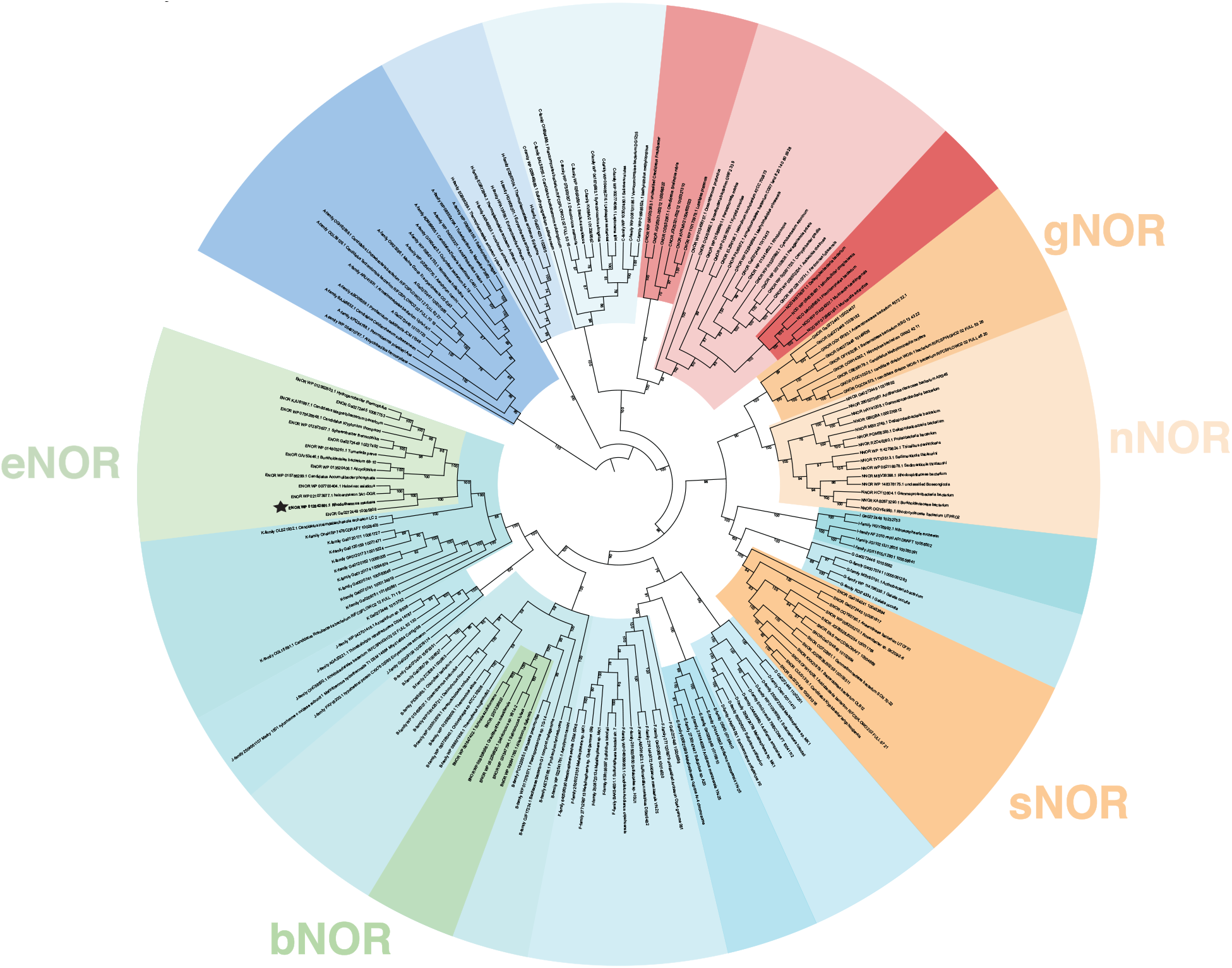
Evolution of nitric oxide reductases. Phylogenetic tree of HCO families. All of the new NOR families are derived from oxygen reductase ancestors. Oxygen reductases are in shades of blue, whereas nitric oxide reductases are in shades of yellow, green and red. The biochemically characterized eNOR from *Rhodothermus marinus* is marked with a black star.

## Biochemical characterization of eNOR

To validate these predictions we isolated and biochemically characterized a member of the eNOR family from *Rhodothermus marinus* DSM 4252, a thermophilic member of the Bacteroidetes phylum. *Rhodothermus marinus* was originally classified as a strict aerobe^23^, but its genome encodes a periplasmic nitrate reductase (NapA), two nitrite reductases (NirS and NirK), and a nitrous oxide reductase (NosZ), suggesting that it may also be capable of denitrification (**Extended Data Figure 1**). Denitrification was not observed under strictly anaerobic conditions, however under microoxic conditions isotopically labeled ^15^NO_3_^-^ was converted to ^30^N_2_ (**Extended Data Figure 2**) demonstrating that *R. marinus* DSM 4252 is capable of complete aerobic denitrification (NO_3_^-^ → N_2_). Blockage of the nitrous oxide reductase with acetylene led to the accumulation of N_2_O (**Figure 3**), suggesting that a nitric oxide reductase was also present in *R. marinus* DSM 4252. No known NORs (cNOR, qNOR, qCu_A_NOR) were found in the genome. However, *R. marinus* DSM 4252 does encode for a member of the eNOR family (**Extended Data Figure 1**).

**Figure 3.**
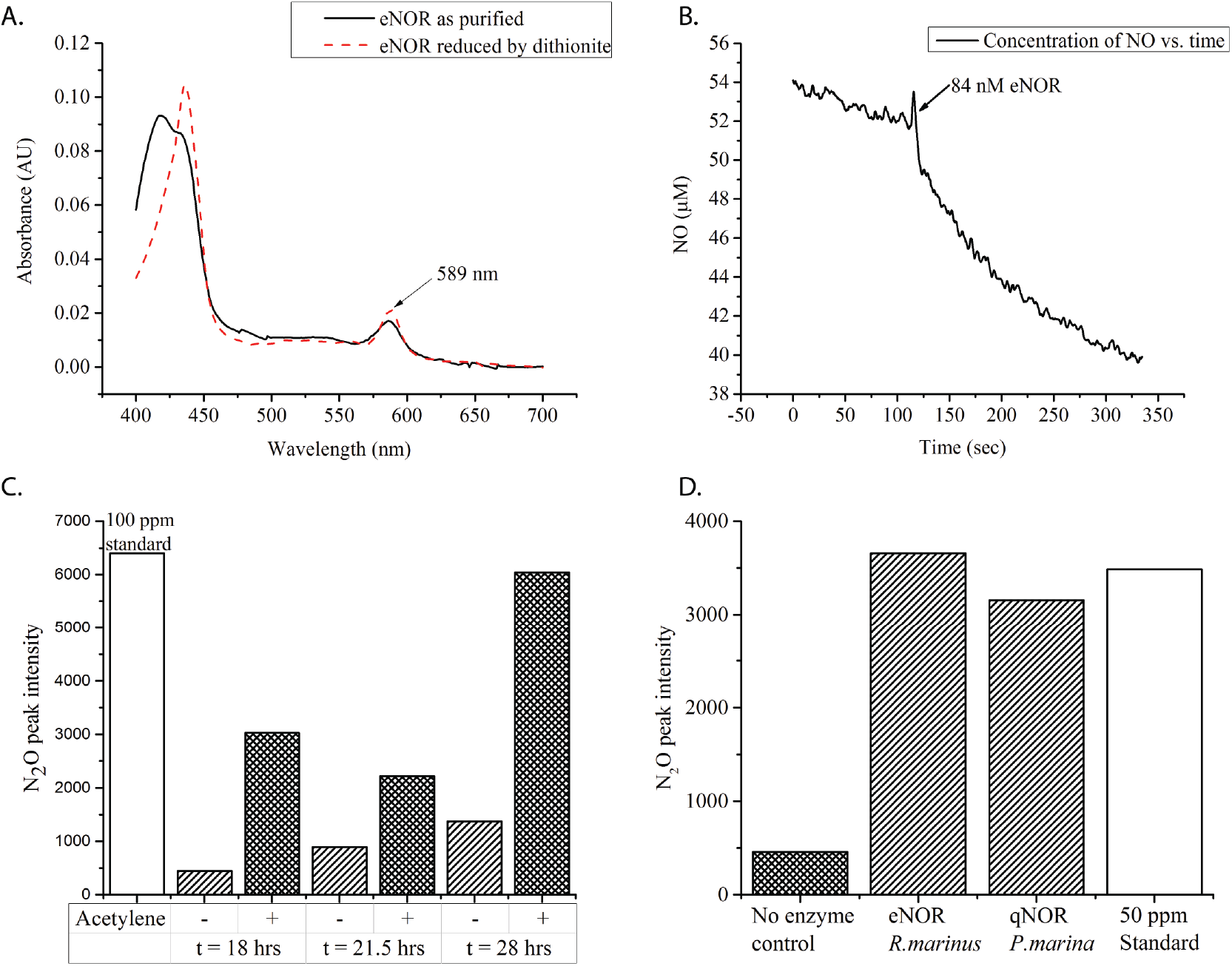
Biochemical Characterization of the eNOR from *Rhodothermus marinus*. a) UV-Vis spectrum of isolated eNOR indicates the presence of an unusual heme *a* signature at 589 nm. b) NO reductase activity was measured with the use of a Clark electrode in the presence of TMPD and ascorbate as electron donor. c) N_2_O accumulation observed in a culture of *Rhodothermus marinus*, in the presence of acetylene d) N_2_O production by eNOR from *R. marinus* and qNOR from *Persephonella marina*.

Isolation and biochemical characterization of the *R. marinus* DSM 4252 eNOR protein verified that it catalyzes NO reduction (at 25°C, k_cat_ = 0.68 ± 0.21 NO.s^-1^ (n = 4)) (**Figure 3**). eNOR was unable to catalyze O_2_ reduction using various electron donors (**Extended Data Figure 3**), clearly showing that it functions solely as a nitric oxide reductase. UV-Vis spectroscopy and heme characterization via mass spectrometry demonstrated that the *R. marinus* DSM 4252 eNOR contains a novel modified heme *a* that is used in both heme sites (**Figure 3 and Extended Data Figures 3 and 4**). A member of the eNOR family was previously isolated from the aerobic denitrifier *Magnetospirillum magnetotacticum* MS-1^16,17^, however its function was never determined. The UV-Vis spectra of the *M. magnetotacticum* eNOR^17^ is identical to the *R. marinus* eNOR, suggesting that the modified heme *a* is a general feature of the family. Mass spectrum analysis of the hemes extracted from eNOR suggest that this heme is A_s_, a previously isolated heme *a* with a hydroxyethylgeranylgeranyl side chain first identified in the B-family oxygen reductase from *Sulfolobus acidocaldarius*^24^. Many eNOR operons contain a CtaA homolog, an O_2_-dependent enzyme that converts heme *o* to heme *a*^25^. This is consistent with the observation that eNOR requires microoxic conditions for expression and suggests that aerobic denitrification might be common in nature.

## Active-site features of novel NORs

In addition to the experimental evidence that both eNOR and bNOR enzymes are NO reductases, there is good evidence that the other newly identified families also perform nitric oxide reduction. The sNOR family has the same active-site structure as the bNOR family, strongly suggesting that it also performs nitric oxide reduction. However, the sNOR and bNOR families are not closely related, providing an example of convergent evolution of active-site structures within the HCO superfamily (**Figures 1 and 2**). Another example of convergent evolution is the nNOR family. It has the same conserved active-site residues as the cNOR and qNOR families (**Figure 1**), but is very distantly related to them (nNOR is related to the B-family, while cNOR and qNOR are related to the C-family). Interestingly, the low-spin heme in nNOR is ligated by a histidine and methionine, which likely raises its redox potential by ~150 mV^26^. This is similar to a modification found in some eNORs, where the low spin heme is ligated by histidine and lysine. The gNOR is the first example of a HCO family that has replaced one of the active-site histidines, residues completely conserved in all other families. A bioinorganic mimic of the gNOR active-site exhibited nitric oxide reduction capability^27^, suggesting that it is likely a functional NOR *in vivo*. The gNOR has a secondary subunit with a cupredoxin fold that is missing the residues required to bind Cu_A_, similar to the quinol oxygen reductase from *E. coli*. Conserved residues that could bind quinol have been identified in gNOR, suggesting that it is a quinol nitric oxide reductase similar to qNOR.

The biochemically characterized (eNOR and bNOR) and proposed (sNOR and gNOR) HCO NORs have active-sites that differ significantly from those utilized by cNOR and qNOR (**Figure 1**). This demonstrates that while oxygen reduction chemistry is constrained to require a redox active tyrosine cofactor, multiple HCO active-site structures are compatible with nitric oxide reduction chemistry. Interestingly, in the currently characterized HCOs Cu_B_ is utilized for O_2_ reduction chemistry and Fe_B_ for NO reduction chemistry. If this pattern is verified for the other predicted NOR families it would suggest that the chemistry performed by HCOs is partially determined by the electronic properties of the active-site metal.

## Bioenergetics of novel denitrification pathways

Although both denitrification and aerobic respiration are highly exergonic processes, most of the enzymes in the denitrification pathway are not coupled to energy conservation, making denitrification significantly less efficient than aerobic respiration^28^. In the HCO oxygen reductases conserved proton channels deliver protons from the cytoplasm to the active-site for chemistry. These same channels are used to pump protons to the periplasmic side. The cNORs and qNORs do not have conserved proton channels from the cytoplasm^15,29^, making them incapable of conserving energy. The eNOR has conserved residues that closely resemble those found in the proton-conducting K-channel within the B-family of oxygen reductases^30^ (**Supplementary Table 1, Extended Data Figure 5**). The sNOR family also has conserved residues in the K-channel region, however they are different than those found in the eNOR and bNOR families. Interestingly, the nNOR family, which has the same active site as cNOR and qNOR, also has a conserved proton channel (**Supplementary Table 1, Extended Data Figure 5**), suggesting that the lack of a proton channel in the cNOR and qNOR may not be due to energy constraints^31^. The conserved proton channels in the eNOR, bNOR, sNOR, and nNOR families would allow them to conserve energy via charge separation, and potentially by proton pumping. Characterization of these new NOR families will be critical for a full understanding of the mechanism of proton pumping in the HCO superfamily, one of the most important unanswered question in bioenergetics.

## Distribution and environmental relevance of NORs

The new HCO NOR families have broad phylogenetic and environmental distributions (**Table 1, Supplementary Tables 2, 3**). The eNOR, sNOR, gNOR, and nNOR families are found in both Bacteria and Archaea. To date, the bNOR family has been found only in the Bacillales order of Firmicutes (**Supplementary Table 2**). Phylogenetic analysis of metagenomic data shows that the majority of eNOR, sNOR, gNOR, and nNOR enzymes are from uncharacterized organisms, suggesting that many more organisms are capable of nitric oxide reduction than previously suspected. Furthermore, the new HCO NOR families are found in a wide variety of environments (**Table 1, Supplementary Table 2**). The sNOR are found in the majority of ammonia oxidizing bacteria sequenced to date and likely play a role in this process. The gNORs are predominantly found in sulfidic environments and may be an adaptation that allows for denitrification in the presence of sulfide, which inhibits other NOR families. The eNOR family is very common in nature, having a broad distribution similar to the cNOR and qNOR families. The eNOR family appears to play key roles in a number of important microbiological processes. They are found in many strains of *Candidatus Accumulibacter phosphatis*, a microbe utilized for phosphate accumulation in wastewater treatment plants during enhanced biological phosphorus removal. The eNOR is highly expressed in transcriptomic datasets from these facilities, demonstrating that *Accumulibacter phosphatis* is capable of complete denitrification *in situ*^32^. eNOR has also been found in microbes capable of performing autotrophic nitrate reduction coupled to Fe(II) oxidation (NRFO). *Gallionellaceae* KS and related strains express an eNOR under denitrifying conditions, suggesting that an individual organism is capable of complete NRFO^33^. eNOR is also commonly found in hypersaline environments (**Supplementary Table 2**) where is might play a role in the adaptation of denitrification to high salt conditions.

**Table 1.** Environmental distribution of the HCO NOR families. Distribution of NOR families in sequenced genomes versus environmental datasets. The newly discovered NOR families account for approximately 2/3 of currently known diversity and 1/2 of the abundance of NORs in nature.

|  | **NCBI-Genomes** | **IMG-metagenomes** | **GTDB-genomes** |
| --- | --- | --- | --- |
| **A-family** | 20290 | 102368 | 45135 |
| **B-family** | 1238 | 4683 | 2021 |
| **C-family** | 13976 | 23015 | 14981 |
| **qNOR** | 4388 | 7680 | 3458 |
| **cNOR** | 2801 | 4824 | 2594 |
| **eNOR** | 68 | 2709 | 547 |
| **sNOR** | 95 | 872 | 344 |
| **bNOR** | 51 | 12 | 200 |
| **nNOR** | 6 | 289 | 32 |
| **gNOR** | 10 | 913 | 156 |
| **NOD** | 8 | 539 | 108 |
| **N2O** | 25 | 597 | n.a. |

Many organisms encode NORs from multiple families (e.g., *Ca*. Methylomirabilis oxyfera has a qNOR, sNOR and gNOR, and *Bacillus azotoformans* has a qNOR, sNOR, and bNOR). This suggests that selection for different enzymatic properties (NO affinity, enzyme kinetics, energy conservation, or sensitivity to inhibitors) or the concentration of O_2_ may be important factors in determining their distribution, similar to what is observed for the HCO oxygen reductase families^8^. Analysis of the presence of denitrification genes (nitrite reductases, nitric oxide reductases, and nitrous oxide reductases) within sequenced genomes indicates that many more organisms are capable of complete denitrification than previously realized. Our current understanding of the diversity of organisms capable of performing denitrification is far from complete.

Our evolutionary analysis shows that nitric oxide reductases have evolved many times independently from oxygen reductases (**Figure 2**). The current data show that NORs have originated from both the B and C-families of oxygen reductases, enzymes that are adapted to low O_2_ environments. These oxygen reductases can reduce NO at high concentrations *in vitro*^34^ so it is not surprising that small evolutionary modifications would lead to enzymes capable of NO reduction at the lower NO concentrations produced during denitrification. The fact that NO reductases are derived from oxygen reductases strongly suggests that complete denitrification evolved after aerobic respiration. This places important constraints on the nitrogen cycle before the rise of oxygen.

## Acknowledgment

We would like to thank NIH for funding this research (Grant #, Principle Investigator: Dr. Robert Gennis). We thank Sylvia Choi for providing pure *ba_3_* oxygen reductase from *Thermus thermophilus* to use as a control for oxygen reductase assays and for heme extraction, Lici Schurig-Briccio for guidance in performing nitric oxide reductase assays with the Clark Electrode and Peter Yau at the University of Illinois’ Mass spectrometric facility for protein identification. We thank Alon Philosof and Connor Skennerton for valuable discussions on bioinformatics analysis. A portion of this research was performed under the Facilities Integrating Collaborations for User Science (FICUS) initiative (award 503546 to R.H.) and used resources at the DOE Joint Genome Institute and the Environmental Molecular Sciences Laboratory, which are DOE Office of Science User Facilities. Both facilities are sponsored by the Office of Biological and Environmental Research and operated under Contract Nos. DE-AC02-05CH11231 (JGI) and DE-AC05-76RL01830 (EMSL).

## List of Tables

**Supplementary Table 1. Putative proton channels in the new NOR families – eNOR, bNOR, sNOR, nNOR.** A list of conserved residues is noted in the table for each family with a reference sequence according to which the residues are numbered. These conserved residues are compared with amino acids found in analogous positions in the B-type oxygen reductase.

**Supplementary Table 2. Distribution of NOR families by phylum in GTDB.** A distribution of all the NOR families within various bacterial and archaeal phyla within the genomes in release 202 of GTDB was analyzed using HMMs that are specific to each NOR family.

**Supplementary Table 3. Distribution of NOR families in various ecosystems as per the IMG database.** A distribution of various NOR families in over 2000 metagenomes on the IMG database was evaluated, and then tabulated according to the environment from which each metagenome is sourced.

**Supplementary Table 4. Denitrification pathways in bacteria and archaea.** An analysis of denitrification pathways in bacterial genomes and archaeal genomes in release 202 of GTDB was performed by searching for the presence and absence of NarGHI, NapAB, NirK, NirS, NosZ, NosD and the NORs in each genome using curated HMMs for each of the proteins.

---

**Extended Data Figure 1:**
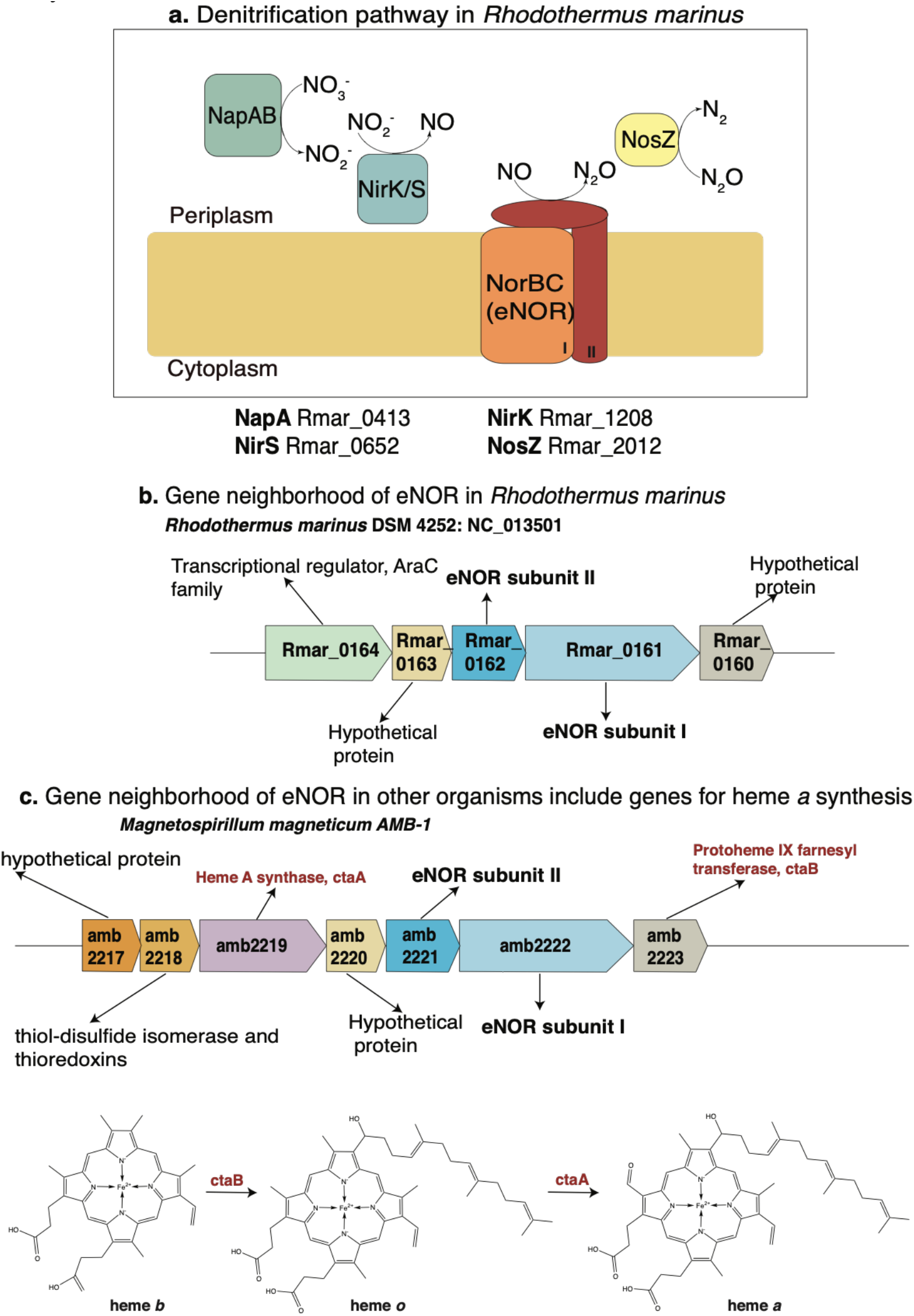
Genome of *R. marinus* encodes for the complete denitrification pathway. a. The genes for NapAB, the periplasmic nitrate reductase (Rmar_0413), nitrite reductases nirK (Rmar_1208) and nirS (Rmar_0652), nitric oxide reductase eNOR(Rmar_0161) and nitrous oxide reductase, nosZ (Rmar_2012) are encoded in the *R. marinus* genome. b. The gene neighborhood of eNOR in *R. marinus*. c. The gene neighborhood of eNOR in *Magnetospirillum magneticum AMB-1* includes ctaA and ctaB, enzymes involved in the biosynthesis of heme *a*.

**Extended Data Figure 2:**
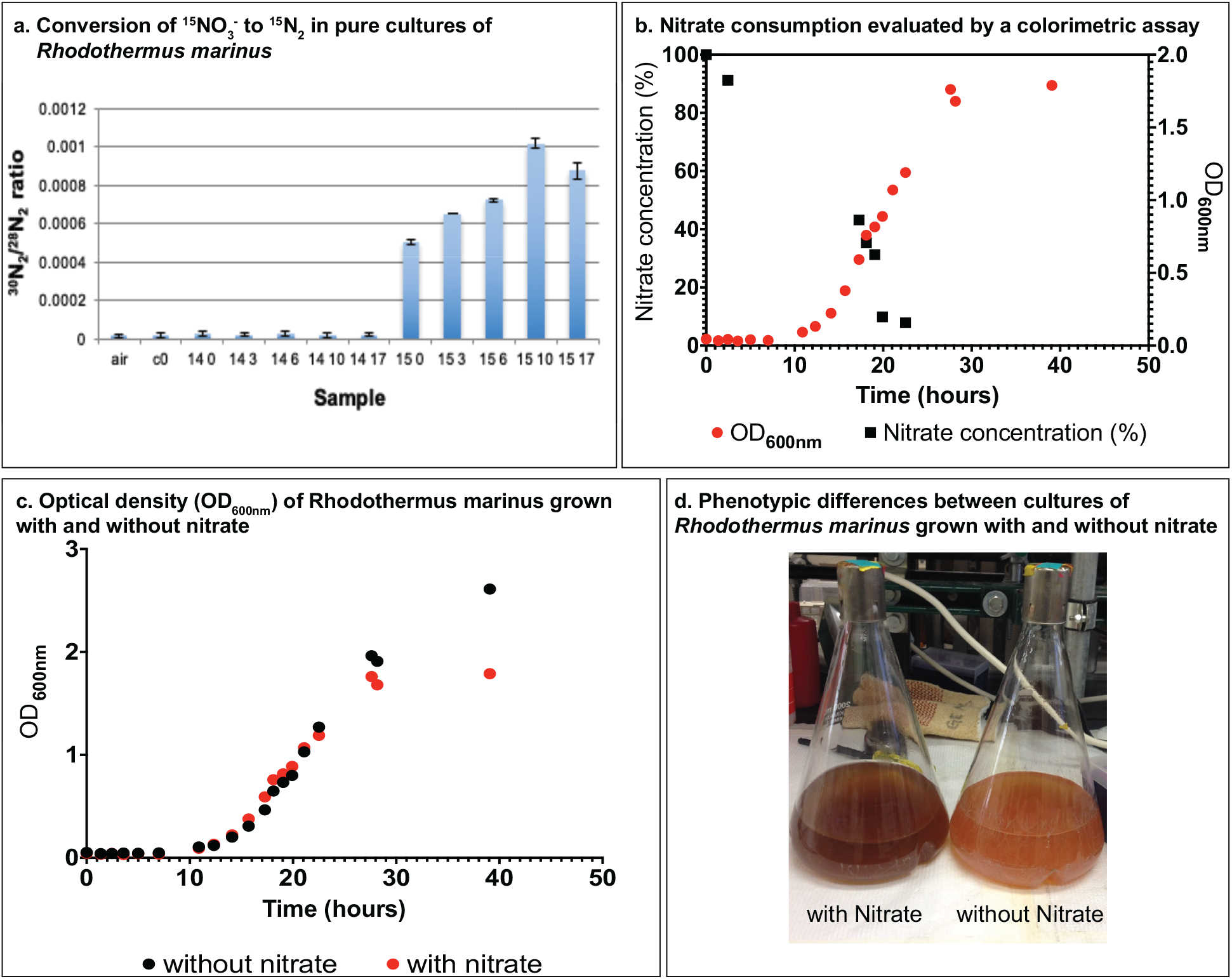
*Rhodothermus marinus* does perform complete denitrification. a. *R. marinus* converts ^15^NO_3_^-^ to ^30^N_2_. Ratio of ^30^N_2_ to ^28^N_2_ for each sample. Air is ambient atmosphere as a standard. C0 is a nitrate-free control. 14 0-17 are cultures grown with unlabeled nitrate, transferred to sealed vials after 0-17 hours, respectively. 15 0-17 are the equivalent samples grown with ^15^N-labeled nitrate. Error bars represent two standard deviations from three replicate GC/MS measurements. ^30^N_2_ enrichments from the ^15^N-labeled samples are over 30-60x higher than background atmospheric ratios, while unlabeled samples have no significant enrichment over background. b. Growth of *R. marinus*, measured using OD_600nm_ over 39 hours. NO_3_^-^ utilization was established by measuring the concentrations of nitrate in the media using a calorimetric assay. c. *R. marinus* growth in rich media was compared under denitrifying and nondenitrifying conditions using OD600nm. d. Phenotypic differences of *R. marinus* cultures, under denitrifying and non-denitrifying conditions.

**Extended Data Figure 3:**
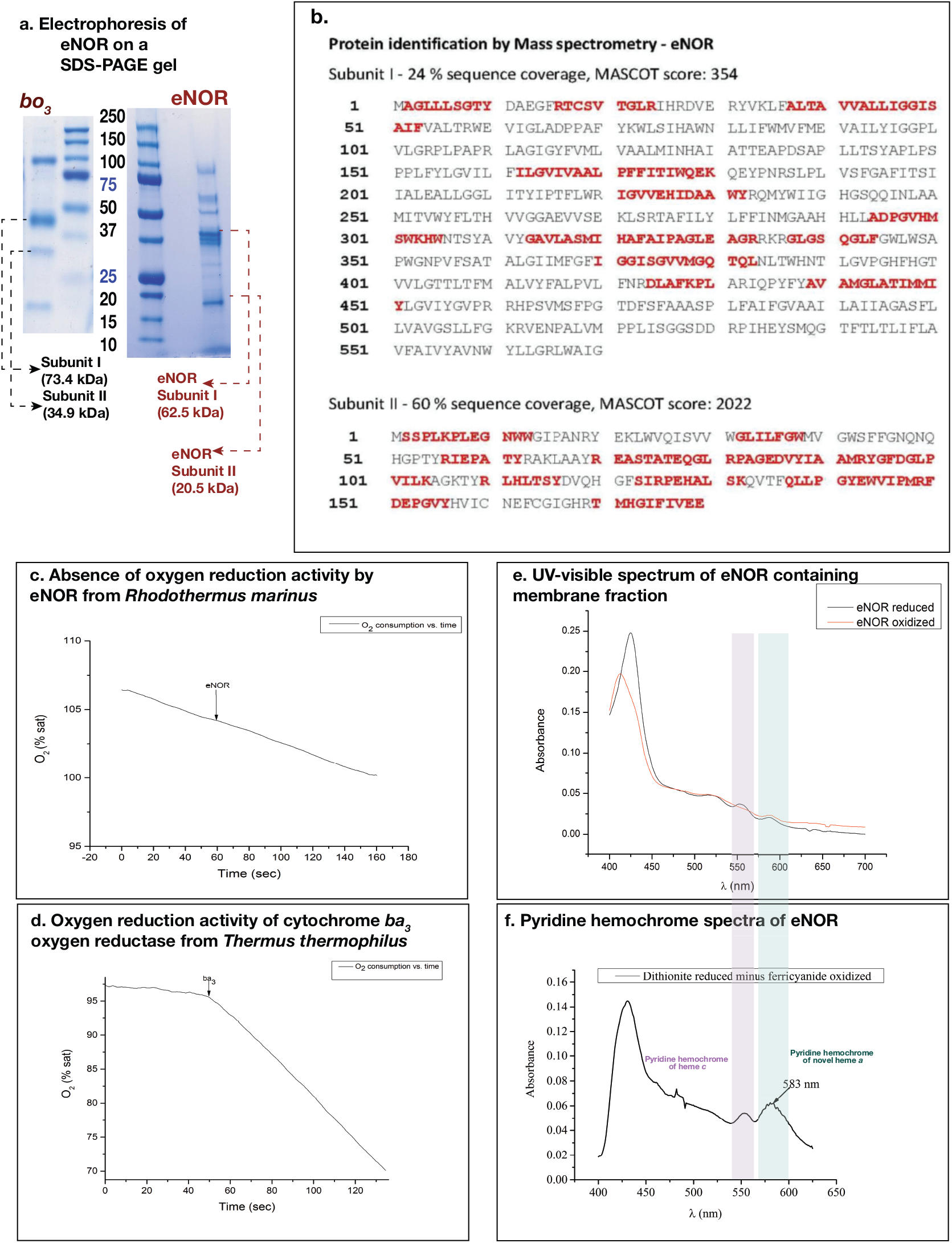
Characteristics of eNOR from *Rhodothermus marinus*. a. SDS-PAGE gel electrophoresis of eNOR shows two bright bands which are estimated to subunits of I and II of the complex. Both subunits appear to run faster than their estimated molecular weight. This is typical for membrane proteins. For comparison, an SDS-PAGE gel of cytochrome *bo_3_* oxidase from *E. coli* is included. b. Mass spectrometric identification of eNOR is confirmed by LC/MS/MS analysis. c,d. Absence of O_2_ reduction by *R. marinus* eNOR, in comparison to robust O_2_ reduction by *T. thermophilus ba_3_*-type oxygen reductase. e. UV-visible spectrum of eNOR f. Pyridine hemochrome-spectra of extracted hemes from eNOR showing a peak which is atypical of hemes *a, b* or *c*.

**Extended Data Figure 4:**
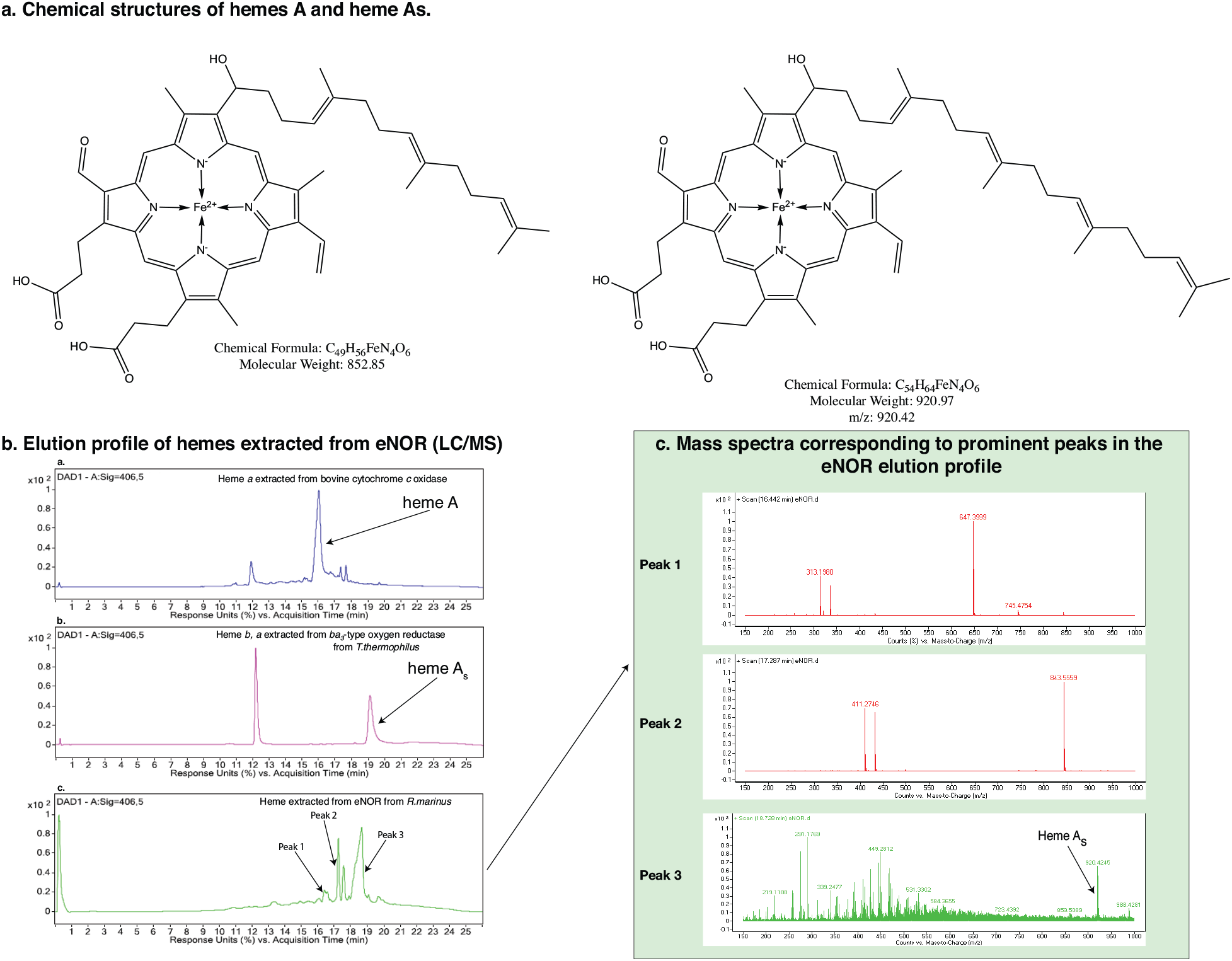
Identification of hemes extracted from eNOR. Comparing the elution profile of extracted hemes from partially purified *R. marinus* eNOR to bovine cytochrome *c* oxidase (A-type, t=16 min), *T. thermophilus ba_3_*-type oxygen reductase (*b*- and A_s_-type hemes, t=12 min and t=19 min) reveals that the heme is most likely an A_s_-type heme. Mass spectra of the peak at ~19 min from the eNOR hemes elution profile confirms that the heme is an A_s_-type heme with a molecular weight of 920 Da.

**Extended Data Figure 5:**
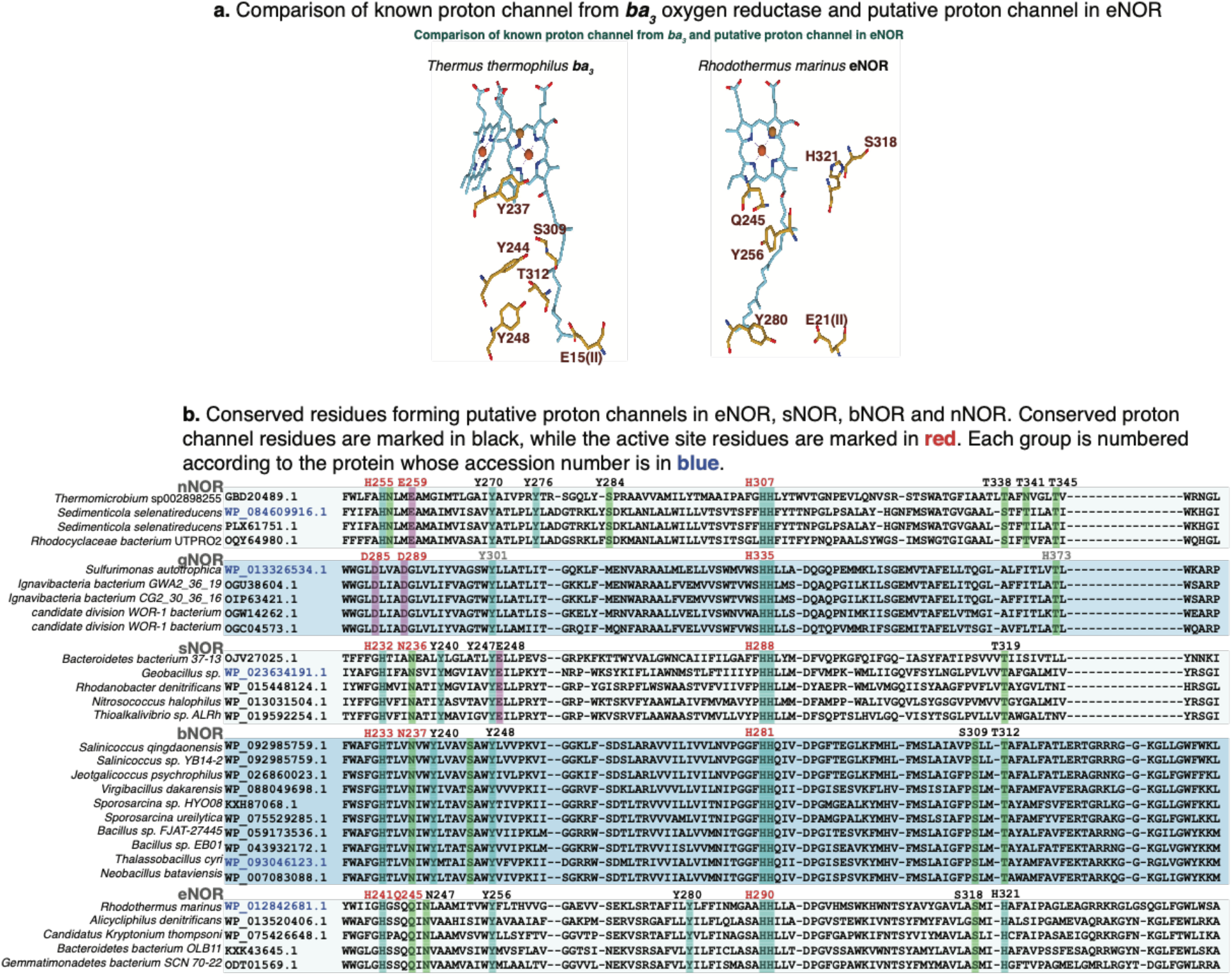
Proton channel in eNOR of *Rhodothermus marinus* and in the NOR families bNOR, sNOR and nNOR. **a.** eNOR contains conserved residues in Helix VII, similar to the location of K-proton channel residues in *T. thermophilus ba_3_*-type oxygen reductase. b. A multiple sequence alignment of the NOR families eNOR, bNOR, sNOR and nNOR show conserved amino acids in analogous location to the K-channel in the B-type oxygen reductase. Some conserved residues are also identified in gNOR and may indicate the presence of a conserved proton channel but they do not map to corresponding residues in the B-type oxygen reductase.

**Extended Data Figure 6:**
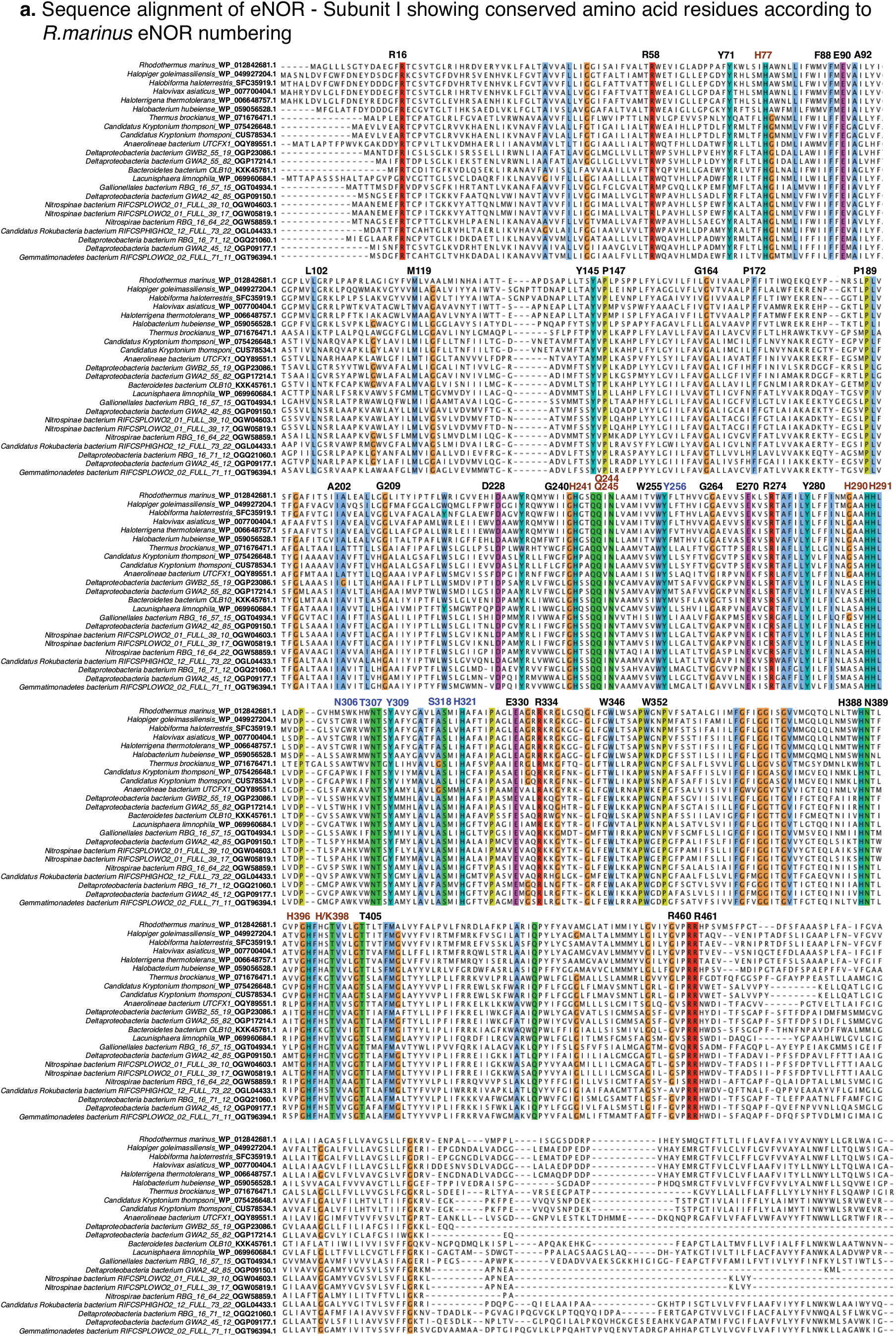
Conserved amino acids in the eNOR family of enzymes. Multiple sequence alignment of 23 eNOR sequences from various taxonomically divergent organisms reveals conserved residues that correspond to the active site ligands, proton channel residues and other sequence features that are unique to eNOR. The active site residues are highlighted in maroon while the proton channel residues are highlighted in blue.

## Materials and Methods

### Growth and Expression Conditions

*Rhodothermus marinus* DSM 4252 was inoculated from frozen stock and grown in 5 ml of DSM Medium 630 with 10 g/L NaCl at 60 °C for 36 hrs. It was then inoculated into a larger secondary culture and grown overnight. 25 ml of the culture was inoculated into 1 L of medium with 30 mM nitrate added. The cells were shaken at 75 rpm and grown at 60 °C. The cells were pelleted by centrifugation at 8000 rpm. The cell pellet was either directly used for protein purification or frozen at −80 °C until the time of use.

### Labeled ^15^NO experiments

We used labeled nitrate (^15^NO_3_^2-^) to verify that *Rhodothermus marinus* DSM 4252 was capable of complete denitrification (NO_3_^-^ to N_2_) using eNOR as the sole nitric oxide reductase. Cultures were inoculated into flasks containing media with either ^14^NO_3_^2-^, ^15^NO_3_^2-^, or no nitrate. The cultures were then allowed to grow microaerobically for a period of time before being subsampled for transfer to sealed vials in order to allow accumulation of gaseous end products. Samples were taken from each media composition after 0, 3, 6, 10, and 17 hours. The headspace was sampled after 20 hours of growth in sealed vials via gastight GC syringe and immediately injected into a Hewlett Packard 5972 gas chromatograph/mass spectrometer. Chromatogram peaks corresponding to isotopologues of NO, N_2_O, and N_2_ were identified by their mass spectra and peak areas were quantified relative to ambient air. As ^15^N cultures were grown in isotopically pure ^15^NO_3_^2-^, complete denitrification should result in accumulation of ^30^N_2_ at a 1:2 ratio relative to nitrate consumption. ^30^N_2_ should only accumulate if eNOR is functioning as part of a complete denitrification pathway. If eNOR does not function effectively as a nitric oxide reductase, then ^15^NO should be seen to accumulate. Instead, only the ^30^N_2_ peak was observed, indicating the eNOR functioned effectively as a nitric oxide reductase for denitrification. Over the course of incubations, ^30^N_2_ was seen to accumulate to more than 50x background. ^14^N samples showed no significant accumulation of ^30^N_2_ above background, confirming that the ^30^N_2_ in ^15^N samples was due to denitrification of labeled nitrate. NO was not seen to accumulate in any of the cultures. These results demonstrate that eNOR is a functional nitric oxide reductase and can be used as part of a complete denitrification pathway.

### Purification of eNOR

The culture of *Rhodothermus marinus*, once harvested, was re-suspended in 100 mM Tris-HCl, pH 8 with 10 mM MgCl_2_ and 50 μg/ml DNase, using a Bamix homogenizer. The resulting solution was spun down at 42000 rpm in a Beckman Ultracentrifuge. The membrane pellet was collected and re-suspended in 20 mM Tris-HCl, pH 7.5, 1 % CHAPS (Affymetrix) to a final concentration of 40-50 mg/ml. The solution was stirred at 4 °C for 1 hr. In this step a lot of peripheral membrane proteins appear to be solubilized and the remaining protein is pelleted by spinning down at 42000 rpm for 1 hr. The remaining pellet is then solubilized in 20 mM Tris-HCl, pH 7.5, 1 % DDM (Affymetrix) at a final protein concentration of around 5-10 mg/ml. The DDM solubilized fraction was once again centrifuged at 42000 rpm to pellet down protein that was not solubilized.

The solubilized protein was then loaded on a DEAE CL-6B (Sigma) column, pre-equilibrated in 20 mM Tris-HCl, pH 7.5, 0.05 % DDM, and subjected to a linear gradient spanning from 0 to 500 mM NaCl. The fraction containing the eNOR, identified using a peak at 591nm, corresponding to the peak of cytochrome ‘a1’ in *Magnetospirillum magnetotacticum*^1^, eluted at around 200 mM salt. This fraction was then loaded on a Q Sepharose High Performance (GE Healthcare) column, pre-equilibrated with 20 mM Tris-HCl, pH 7,5, 0.05 % DDM and then eluted in a gradient from 0 to 1 M NaCl. The eNOR containing fraction was eluted at around 250 mM salt and the eluted fraction was then loaded on a Chelating Sepharose (GE Healthcare) column, loaded with Cu^2+^ and equilibriated with 20 mM Tris-HCl, 500 mM NaCl, as previously described for cytochrome *caa_3_* from *Rhodothermus marinus*^2^. The eNOR fraction was once again loaded on a Q Sepharose High performance column, and a gradient was run between 0 and 300 mM NaCl at low flow rates (0.5 ml/min) and the first peak was found to be the eNOR.

### Gel Electrophoresis

The purified eNOR was run on a Tris-Hepes 4-20 % acrylamide gel (NuSep) in the recommended Tris-Hepes-SDS running buffer at 120 V for ~1 hr. The protein was visualized and compared to the Precision Plus Protein™ Dual Color Standards (BIO-RAD).

### UV-Visible Spectroscopy

All spectra were recorded on a HP Agilent 8453 UV-Vis spectrophotometer using a quartz cuvette from Starna Cells (No. 16.4-Q-10/Z15). Potassium Ferricyanide was used to obtain the oxidized spectrum, and dithionite was used to obtain the reduced spectrum.

### Pyridine Hemochrome Assay

The hemes in eNOR were analyzed using a pyridine hemochrome assay^3^. A stock solution of 200 mM NaOH with 40 % pyridine was prepared. The stock solution was mixed 1:1 with the protein and an oxidized spectrum was obtained by adding 3 μl of 100 mM K_3_Fe(CN)_6_. A reduced spectrum was similarly prepared by adding a few crystals of sodium dithionite. The reduced minus oxidized spectrum was used to identify the heme co-factors

### Heme extraction and HPLC Analysis

The hemes from eNOR were extracted and analyzed using an HPLC elution profile according to established protocols^4,5^. 50 μl of eNOR was mixed with 0.45 ml of acetone / HCl (19:1) and incubated for 20 minutes at room temperature after shaking. The mixture was centrifuged at 14,000 rpm for 2 minutes, followed by addition of 1 ml of ice cold water, and 0.3 ml of 100% ethyl acetate to the supernatant. The water/ethyl acetate mixture was vortexed and centrifuged again for 2 minutes. The ethyl acetate phase was recovered and concentrated using a speed vac.

The extracted hemes were analyzed using an Agilent 1290 Infinity LC attached to an Agilent 6230 TOF LC/MS equipment by separation using an Agilent Eclipse Plus C18 column (2.1×300 mm, 1.8 μm, 600 bar) and an acetonitrile (0.05 %TFA) / water (0.05 % TFA) gradient from 20 to 95 %.

### NO reductase activity verification using GC

Anaerobic reaction conditions were set up in a 5 mL clear serum vial (Voigt Global Distribution, Inc) sealed with a 20 mm rubber stopper, by passing N_2_ through 2 ml of 20 mM KPi, 0.05 % DDM, pH 7.5 with 1 mM TMPD, 5 mM Ascorbate. A control was performed by adding only 50 μM NO. Sample reactions were begun by adding eNOR to a final concentration of 100 nM. The reaction was incubated at 42 °C for half an hour before the headspace was injected into an HP Agilent 5890 Series GC, fitted with a TCD and ECD (SRI Instruments) for verification of N_2_O production.

### Turnover measurement using a Clark electrode

A sealed chamber fitted with an ISO-NO (World Precision Instruments) electrode was used for NO reductase activity measurements. 1 mM TMPD or 100 μM PMS and 4 mM Ascorbate were was added to 2 ml 50 mM Citrate, pH 6, 0.05 % DDM in the reaction chamber and all traces of oxygen were removed by passing water-saturated Argon for 20 minutes through the solution. This is similar to the protocol described for cNOR from *Thermus thermophilus*^6^. The buffer system also contained an oxygen scavenging system constituting 100 nM catalase, 35 nM Glucose oxidase and 90 nM Glucose. The NO reduction traces were recorded using a Duo-18 (World Precision Instruments), and activities calculated from the slope of the traces.

### LC/MS/MS analysis

Mass spectrometric analysis was conducted at the Protein Sciences Facility, Roy J Carver Biotechnology Center, University of Illinois, Urbana, IL 61801 using a Thermo LTQ Velos ETD pro mass spectrometer. For liquid samples, the samples were cleaned up using G-Biosciences Perfect Focus (St. Louis MO) prior to digestion with trypsin. Digestion was done using proteomics grade trypsin 1:20 (G-Biosciences, St. Louis, MO) and a CEM Discover Microwave Reactor (Mathews, NC) for 15 minutes at 55° C at 50 Watts. Digested peptides were extracted 3X using 50% acetonitrile containing 5% formic acid, pooled and dried using a Speedvac (Thermo Scientific). The dried peptides were suspended in 5% acetonitrile containing 0.1% formic acid and applied to LC/MS.

HPLC for the trypsin digested peptides was performed with a Thermo Fisher Dionex 3000 RSLCnano using Thermo Acclaim PepMap RSLC column (75 μm x 15 cm C-18, 2 μm, 100Å) and a Thermo Acclaim PepMap 100 Guard column (100 μm x 2 cm, C-18, 5 μm, 100Å), solvents were water containing 0.1% formic acid (A) and acetonitrile containing 0.1% formic acid (B) at a flow rate of 300 nanoliters per minute at 40 °C. Gradient was from 100% A to 60% B in 60 minutes. The effluent from the UHPLC was infused directly into a Thermo LTQ Velos ETD Pro mass spectrometer.

Control and data acquisition of the mass spectrometer was done using Xcalibur 2.2 under data dependent acquisition mode, after an initial full scan, the top five most intense ions were subjected to MS/MS fragmentation by collision induced dissociation. The raw data were processed by Mascot Distiller (Matrix Sciences, London, UK) and then by Mascot version 2.4. The result was searched against NCBI NR Protein database.

### Analysis of heme-copper oxygen reductase phylogeny and distribution in environmental datasets

We performed a large-scale analysis of heme-copper oxygen reductase (HCO) protein sequences in the NCBI and IMG databases with BLASTP using an e-value of 1e-^3^ to generate a database of HCO sequences that had at least some of the conserved amino acids previously identified in subunit I^7^. We then used the database of HCOs, filtered it with a sequence cut-off of 50% to generate the multiple sequence alignment, **MSA1**. A phylogenetic tree (**Figure 2**) was inferred using IQ-TREE 2^8^ with the Dayhoff substitution matrix, Gamma model of rate heterogeneity and 1000 ultrafast bootstraps. Using the curated HMMs for each of the HCO family oxygen reductases^9^, we probed release 202 of the Genome Taxonomy Database^10^ for distribution of the NOR families – eNOR, bNOR, sNOR, nNOR, gNOR, cNOR and qNOR – in bacteria and archaea. Curated HMMs for the nitrate reductases (NapAB, NarGH), nitrite reductases (NirK, NirS) and nitrous oxide reductases (NosD and NosZ) were sourced from the HMMs database of MagicLamp^11^.

Analysis of HCO distribution in various ecosystems were performed using the metagenomes in the IMG database. Approximately 2300 metagenomes were identified which were sourced from 44 environments identified by IMG. The number of different HCOs in each of these environments were extracted using BLASTP and query sequences that belong to each of the different HCO families.

